# Opponent-flipping tactics in aggressive interactions of the cricket *Gryllus bimaculatus*

**DOI:** 10.1101/2025.11.13.688168

**Authors:** Akihisa Murata, Kanako Takemoto, Hitoshi Aonuma

## Abstract

Male crickets engage in intense aggressive behavior, competing for resources. In this study, we focus on the quick movements during tactile combat in the cricket fight, to understand how they defeat the opponents. We performed kinematic analysis following high-speed cam recording of the fight. High-speed cam recordings showed that the attacker jumped to the head of the attacked cricket and thrusted it backwards. The attacked cricket was sometimes flipped over and tended to retreat. To understand how the attacker jumps effectively to flip over the opponent, we compared the attack-jump and escape-jump. The kinematics analysis demonstrated that the attack motion is different from the jump in the case of escaping from threats. The attacker cricket adjusted the direction of its body using its forelegs. The mandibles were used to hook onto the head of the attacked cricket. The attacked cricket moved its hindlegs with different kinematics to jump in the case of escape and exerted greater velocity. These findings advance our knowledge of how animals utilize their body depending on the situation.

## Introduction

Most animals fight between conspecifics for resources such as food, territory, and mates (Lorenz, 1963). To win the fight, the tactics are crucial because the results are dependent on the actions of both an animal and its opponent (Blanchard and Blanchard, 1977; Siva-Jothy, 1987). Understanding what tactics an animal uses is important for understanding its aggressive behavior.

To carry out the tactics effectively, it is necessary for animals to utilize their own body. Animals attack their opponents with unique movements that are dependent on their morphology. For instance, the movements of rhinoceros beetles when throwing an opponent differ depending on the species-specific shape of their horns (McCullough et al., 2014). Hermit crabs, *Pagurus bernhardus*, fight by rapping each other’s shells, and the winner impacts the opponent more effectively (Briffa and Fortescue, 2017). Examining the tactics used by different animals leads to a better understanding of how they move and utilize their physical characteristics.

Male crickets are known for exhibiting aggressive fighting behavior (Alexander, 1961). Each action in the fight has been categorized. When they encounter a conspecific male, crickets exhibit a behavior called antenna fencing, in which they strike their opponents with their antennae like a whip to assess an opponent’s motivation (Hack, 1997; Rillich et al., 2007). If neither cricket retreats during fencing, they finally escalate to physical combat (Adamo and Hoy, 1995; Alexander, 1961; Stevenson et al., 2000). However, the details of the movements in the combat have remained unclear, because it is difficult to observe the aggressive behavior of crickets, which happens quickly, and because they do not possess any obvious weapons. In this study, we addressed how crickets (*Gryllus bimaculatus*) defeat their opponent.

For a better understanding of the tactics in fighting, it is necessary to have information about the motion and the role of different parts of the body. It is reported that fighting crickets display an aggressive behavior in which they jump using their hindlegs to thrust their opponent away (Adamo and Hoy, 1995). Indeed, the well-developed hindlegs are one of the most distinctive features of the Orthoptera, including crickets. Some studies have reported that crickets bite their opponents when in combat (Alexander, 1961; Stevenson et al., 2000). The cricket *Gryllus pennsylvanicus* exhibits sexual dimorphism in its mandibles (Judge and Bonanno, 2008), suggesting that the mandibles may be used for fighting. Forelegs are sometimes used for fighting in other insects (Chen et al., 2002), but the role of the forelegs in the attack behavior of crickets is not yet understood. We therefore investigated how these body parts are utilized for attack.

Here, we report the detailed motion of the attack of crickets. We recorded cricket fighting using high-speed cameras and performed kinematic analyses. Because we predicted that the jump during attack would differ from that during escape, we compared jumping during an attack with jumping during an escape. We found that crickets move their forelegs, hindlegs, and head in characteristic ways when jumping in attack, which are distinct from the ways they move during escape jumps.

## Materials and Methods Animals

*Gryllus bimaculatus* (De Geer) used in this study were raised in a laboratory colony. They were reared under a 12 h: 12 h light: dark cycle (lights on at 6:00 h) at 28 ± 2℃ in plastic cases (28.5 cm × 40.7 cm × 18.5 cm). They were fed a diet of insect food (Sankyo Lab, Tokyo, Japan) and provided with water *ad libitum*. Adult males that had molted within the previous 1 week were randomly selected for this study. To increase aggressive interactions, each cricket was kept isolated in a 100 ml glass beaker for 3 days prior to behavior experiments. During isolation, they were fed chopped carrots.

### Behavior experiments

Behavior experiments were performed between 12:00 h and 18:00 h. Paired crickets were paired for body size. The body mass of the crickets used in this study was 0.65 ± 0.15 g (mean ± SD; Suppl. 1), and the mass difference within the pair did not exceed 0.05 g. To observe the fighting behavior, a pair of crickets was introduced into a test arena (made of acrylic, φ = 200 mm, height 100 mm) whose floor was covered with a matte silicon sheet. The mat was washed to remove the smell of the crickets after each experiment. They were initially separated from each other by an acrylic partition in the arena to prevent any contact and to calm the animals prior to experimentation. At the start of the experiment, the partition was slowly removed, and the crickets’ behavior was observed and recorded until a dominance hierarchy within the pair was established. We focused on tactile combat escalation between males. They were recorded using two synchronized cameras high-speed cam system (HAS-U2M, Ditect, Tokyo, Japan) at 2,000 frame sec^-1^ and a shutter speed of 200 µs.

We compared the motion between the attack jump and the escape jump. The escape jump was induced by an air puff stimulation using an air compressor (LM-8000, Lutron, Pennsylvania, USA). The stimulation was supplied 30 cm away from the cricket through an air tube with a fine nozzle (φ = 1 mm). The velocity of the airflow was 2.5 m _s_^-1^_._

### Kinematics analysis

The high-speed movies of the cricket behavior were used for kinematics analysis. We performed tracking of each point of interest for each video using DeepLabCut (Mathis et al., 2018) with a computer (Ubuntu 20.04.4 LTS, CPU: Intel Core i7-8700K, Intel, California, USA, GPU: Geforce GTX 1080 Ti, NVIDIA, California, USA × 2). Then, we constructed the 3D trajectory of each trial using 3D DeepLabCut (Nath et al., 2019). For the video of the fighting, a total of 418 images were manually labeled with 15 points for each insect (Suppl 1); the base of the antenna, mandible, the joint between head and thorax, the joint between thorax and abdomen, the joint between coxa and femur, the joint between femur and tibia, and the joint between tibia and tarsus. For the escape behavior videos, a total of 553 images were similarly labeled as another dataset. Points were labeled on the side facing the camera, either left or right. Using labeled images, the neural network was trained for 500,000 iterations.

The angle of the longitudinal body axis to the ground (*θ*_Axis_) and the angle of the front side of the head to the body axis (*θ*_Head_) were measured using the open-source software Kinovea 0.9.5 (https://www.kinovea.org/). The angle *θ*_Axis_ was measured as the angle between the line from the base of the antennae to the posterior end of the abdomen and the ground in the frame when the hindlegs of the jumping individual leave the ground. The angle *θ*_Head_ was measured as the angle between a straight line tracing the dorsal side of the mandible and a straight line tracing the dorsal side of the thorax before and during the jump.

The initial velocity and acceleration were calculated based on the three-dimensional coordinates obtained by tracking. For the data of each video, we selected a head point or a thorax point and numerically differentiated the displacement between the frame where the hind legs start to extend and the frame where the hind legs leave the ground.

Each angle was calculated from the dot product of spatial vectors as shown in the following equation:

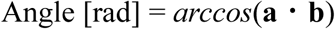

**a** and **b** represent the spatial vector connecting two nodes of the coarse-grained body. To remove noise, the data were filtered by a simple moving average with 9 frames.

## Statistical Analysis

Statistics were calculated using R version 4.2.2 (R Development Core Team, http://www.R-project.org) on RStudio (2023.06.1 Build 524, RStudio PBC, Boston, USA). We used Student’s t-test for a comparison between two groups. To calculate the mean and deviation of circular data, we used the functions ‘mean.circular’ and ‘angular.deviation’ from the R package ‘circular’. For comparison between two groups of circular data, we performed the Mardia-Watson-Wheeler test using the function ‘watson.wheeler.test’ (r package ‘circular’). We applied the Mann-Whitney U test for comparison between two groups of the number of times.

To obtain representative trajectories of legs in each jump, we constructed a statistical model based on the method in (Aonuma et al., 2023). Its parameters were estimated by using the probabilistic programming language Stan through the R package “rstan”. Our model is a state-space model consisting of a first-order trend term. The formula for the Angle at each time point was given by:

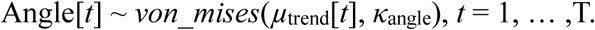

This formula contains three parameters: *t*, *µ*_trend_, and *κ*_angle_, where *t* represents time steps. *κ*_angle_ value represents the concentration of the von Mises distribution and is determined as the overall value through each timestep, and the *µ*_trend_ value constrains the current angle value to be similar to the previous value. It was given by:

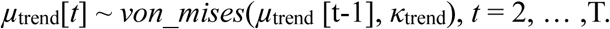

*κ*_trend_ value is also the concentration determined by the overall timestep, the same as *κ*_angle_.

Sampling was performed using the Markov chain Monte Carlo method (MCMC) with 5,000 iterations and 4 chains from experimental data. Iterations 2,501 to 5,000 were sampled. The convergence of sampling was confirmed by drawing a trace plot.

## Results

### Fighting between males

A cricket showed antennal fencing and body jerking when it encountered a conspecific male. If the opponent did not retreat from the fight, the cricket opened its mandibles and quickly attacked the opponent. At the beginning of the tactile combat, the attacking cricket showed boxing-like behavior (Movie 1). In this phase, the attacker cricket used its forelegs to step on the opponent’s foreleg (Movie 1). This boxing behavior is similar to that shown in fruit fly fighting (Chen et al., 2002), but unlike flies, crickets did not stand on their hind legs alone. The defending cricket also showed boxing behavior. Therefore, this boxing behavior appears to be an action to create distance between animals during an attack. During this boxing phase, the mandibles of the attacking cricket were kept fully open (Fig. 2). After the boxing, the cricket placed its open mandibles beneath the mandibles of the defending animal (Movie 2). Then the attacking cricket leaned forward and jumped upwards, which made the defending cricket flip over (Fig. 1). We call this hindleg jump, made by the attacking cricket, an attack jump.

**Fig. 1.**
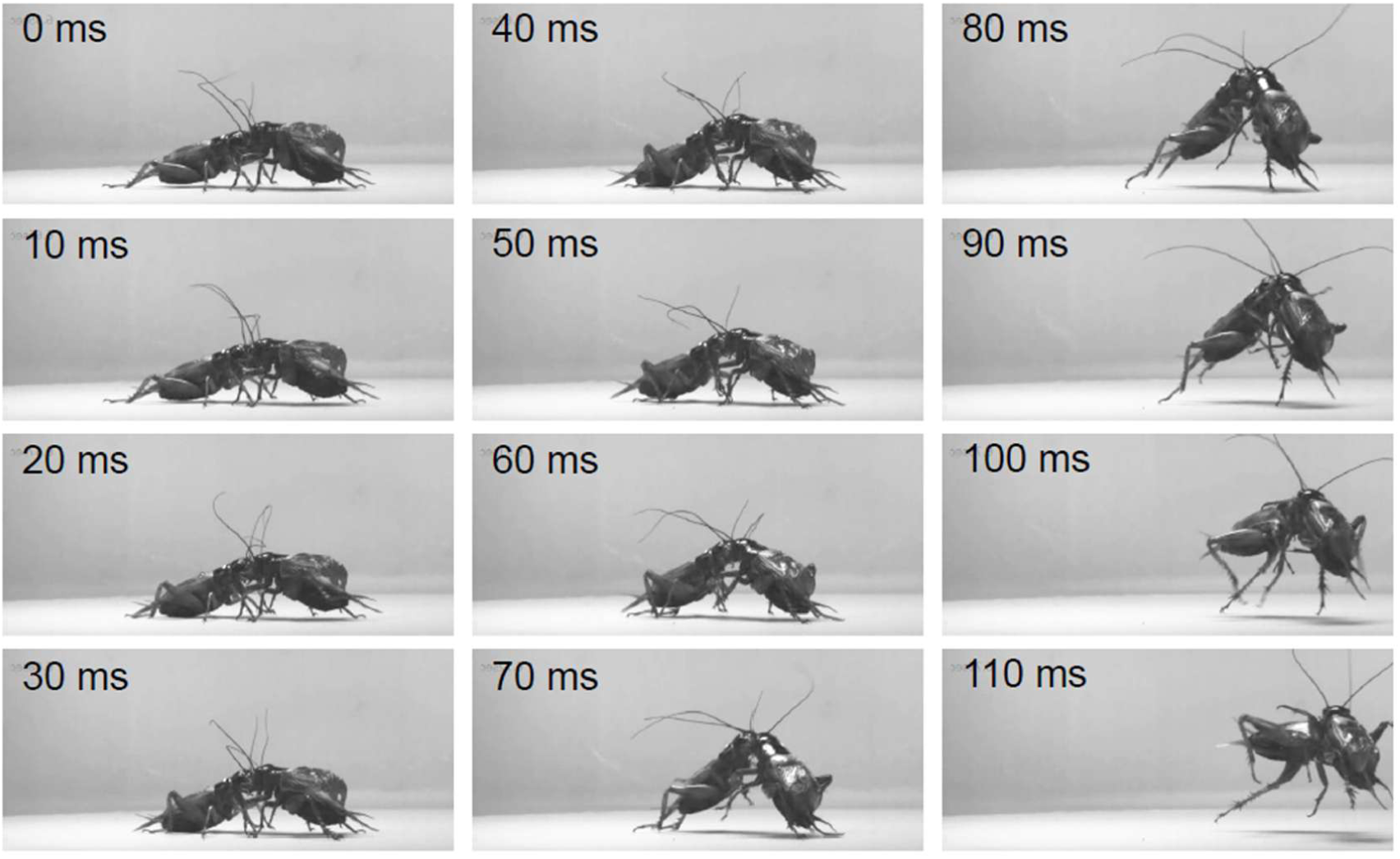
Sequential snapshots of the fighting. The cricket on the left side flipped over the opponent while jumping. The high-speed movie was recorded at a frame rate of 2,000 fps.

**Fig. 2.**
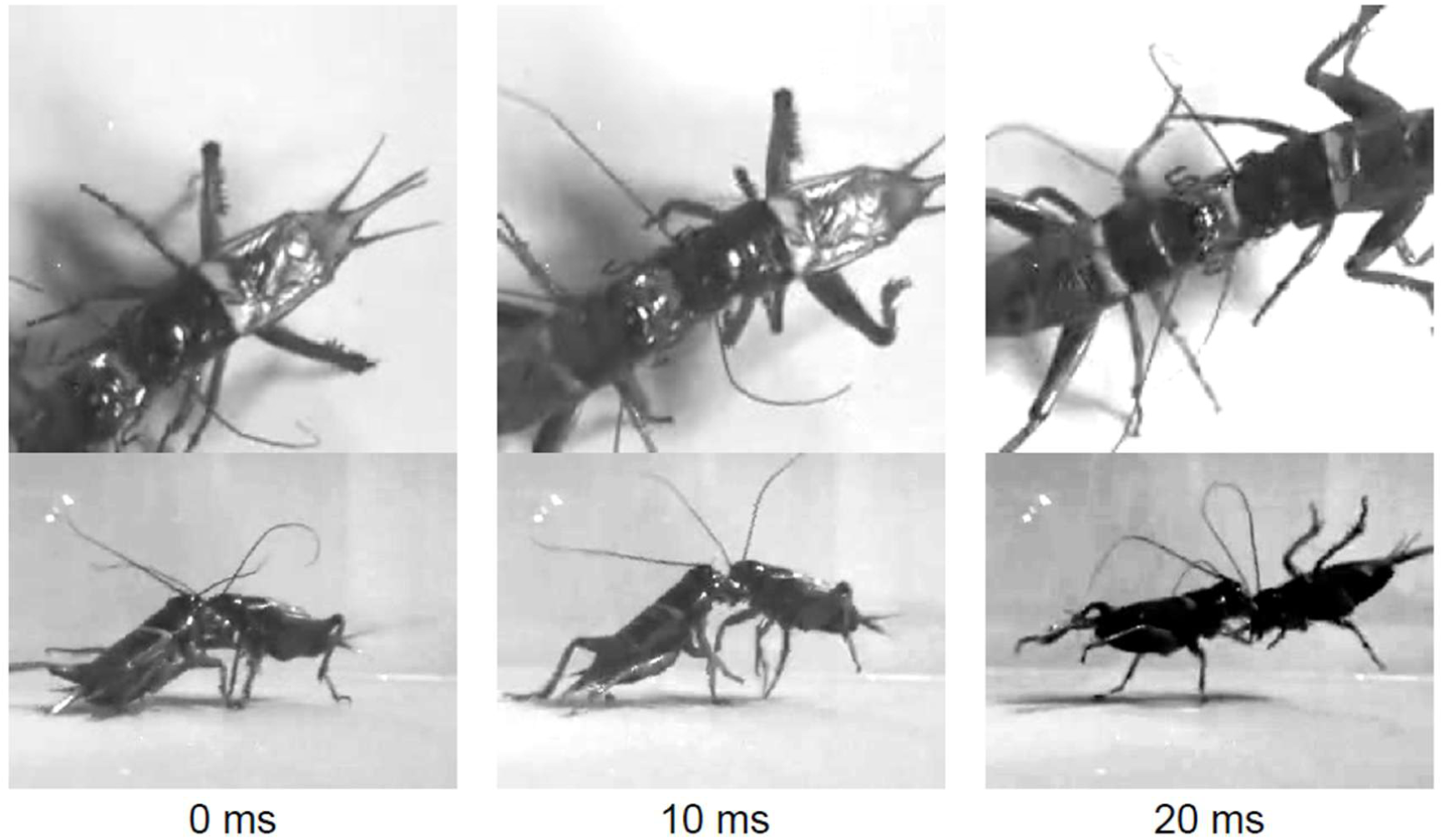
Opened mandibles during attack. Crickets spread their mandibles during the fight. Fights were recorded vertically and horizontally using a high-speed camera with a frame rate of 500 frames s^-1^. The mandibles of the attacker were traced with a dotted line.

In 36 out of the 37 pairs, the attacking crickets kept their mandibles open during the mandible engagement phase and the flip. Only one attacking cricket bit the attacked cricket. This happened when the body of the opponent accidentally touched the mandibles. These results indicate that attacking cricket aims to flip over the opponent to win the fight. We also examined how many times attacks resulted in successful flips and how many times flips occurred during a single fight (Fig. 3). In total, 29 of 38 pairs performed an intense fight. The pairs that did not show an intense fight were excluded from the count. In many cases, the fight ceased when the cricket was flipped over once (Fig. 3). An attack jump would thus be effective in making the opponent give up the fight.

**Fig. 3.**
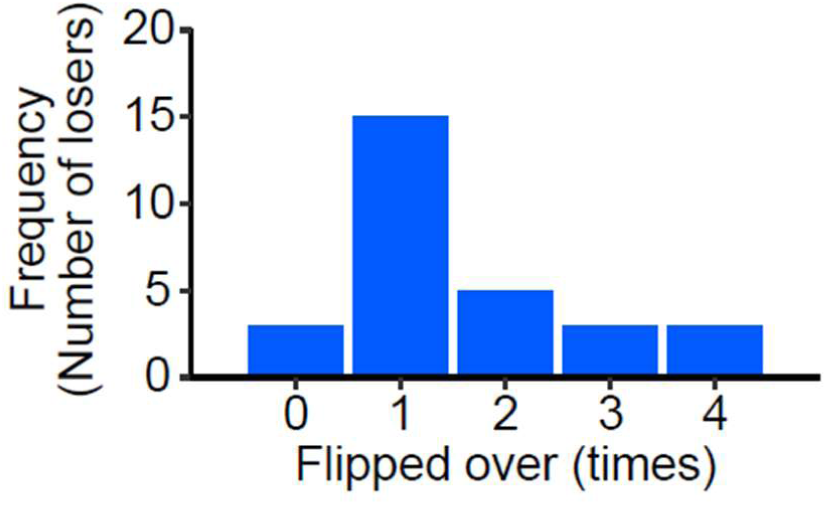
The Frequency with which losers were flipped over in fighting. Crickets tend to lose when they are flipped over only once. In total, 29 pairs performed intense fights among all observed (38 pairs). The pairs that did not show an intense fight are excluded from the count.

The attack jump can be categorized into 2 phases: the preparation phase and the push-off phase. In the preparation phase, the attacking cricket lowered its abdomen to the ground and flexed both hindlegs. Then, the attacker thrust out his mandibles and hooked them onto the opponent’s mandibles. Throughout this phase, both animals extended forelegs and midlegs to support the lifted body. (2) In the push-off phase, the attacker jumped and pushed the opponent with his mandible hooked on the opponent’s mandibles. The attacker’s hindlegs were fully extended, and all legs were used to kick off the ground.

### Kinematics analysis of hindlegs in jumping

To investigate the effectiveness of the attack jump, we performed kinematics analysis and compared it with the escape jump. For attack jumps, we filmed a total of 82 pairs and analyzed 10 sets of videos that were successfully captured. We then filmed a total of 44 escape jumps, and 10 sets of videos that were successfully captured and selected from them were analyzed.

Hindlegs mainly contribute to generating the power for jumping. To confirm whether *G. bimaculatus* changes the motion of its hindlegs to attack, we analyzed the kinematics of the femorotibial joint using 3D tracking (Fig. S1). Crickets took longer to prepare for the attack jumps, and extended their hindlegs further for them, than for escape jumps.

In the case of attack jumps, the leg joints moved in a stereotypical pattern regardless of the initial posture. Crickets flexed their hindlegs prior to jumping, and then fully extended them. As mentioned above, we separately analyzed the movement separated into two phases. We defined the preparation phase as the stage before crickets began to extend their hindlegs (–40 ∼ 0 ms), and the push-off phase as the phase when crickets were leaping (0 ∼ 40 ms). The time at which the angle of the femorotibial joint started to open was designated as *t* = 0. In the preparation phase, the angle of the femorotibial joint of the hindleg (*θ*_hFT_) decreased according to flexion of the hindlegs (Fig. 4A). The angle *θ*_hFT_ was initially in the range of 73 deg to 129 deg (at -40 ms) and decreased by about 30 degrees for each insect. Then, this angle was maintained for about 40 ms. In the statistical model, the angle *θ*_hFT_ decreased from 96 deg to 68 deg as shown in the 50-percentile curve (hereafter referred to as the estimated curve).

**Fig. 4.**
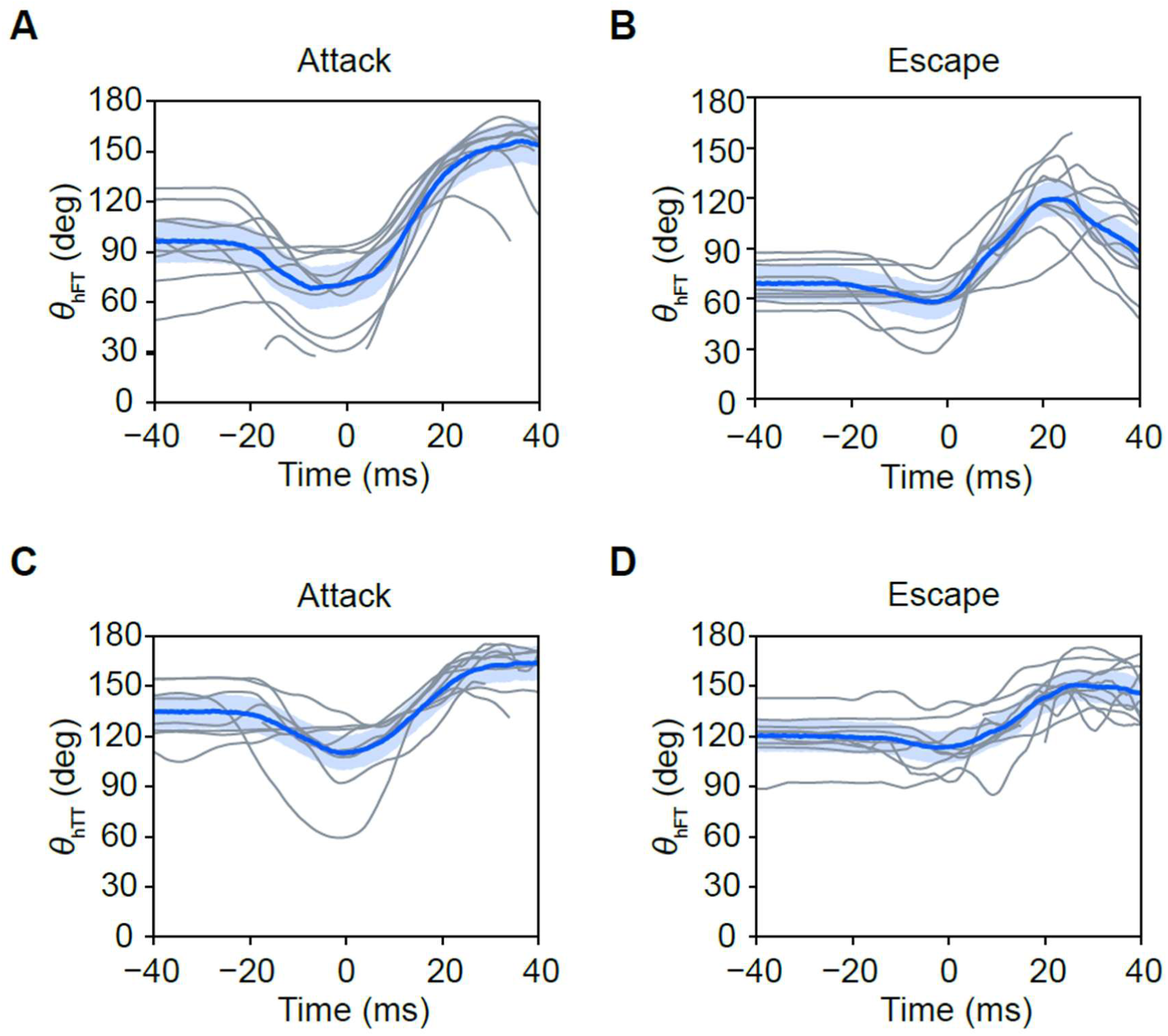
Time validation of the angle of the hindleg during attack and escape jumps. (A, B) Time validation of the angle of the femorotibial joint of the hindleg (*θ*_hFT_) in attack and escape jumps. (C, D) Time validation of the angle of the tibiotarsal joint of the hindleg (*θ*_hTT_) in attack and escape jumps. The angles were calculated for each frame from 3D coordinates. All data series were aligned with a time point when crickets start to extend their hindlegs as 0 ms. Colored lines indicate the estimated 50th percentile, and the gray area indicates the estimated 25th–75th percentile that was based on the statistical model.

In the push-off phase, the angle *θ*_hFT_ greatly increased (by about 90 deg) in about 30 ms and reached a maximum of 123 deg to 171 deg. There was little variation in this motion between individuals. The angle *θ*_hFT_ of the estimated curve increased from about 70 deg to 160 deg in 30 ms. In the case of the escape jump (Fig. 4B), some crickets flexed their hindlegs prior to jumping, but others barely flexed them. Accordingly, the angle *θ*_hFT_ barely decreased in the preparation phase, from 83 deg to 67 deg on the estimated curve. In the push-off phase, the angle *θ*_hFT_ increased by about 60 deg in 20 ms and reached a maximum of 103∼159 deg. The estimated curve indicated that the angle *θ*_hFT_ increased from about 70 deg to 130 deg in 20 ms. This change was smaller, and the time spent jumping was shorter than that of attack jumps.

We then analyzed the kinematics of the tibiotarsal joint because the tarsus is also a component of the mechanical system of the leg. It is known that the movement of the tarsus is linked to that of the tibia in locusts, which belong to the same order (Orthoptera) as crickets (Burrows and Horridge, 1974; Meruelo et al., 2014). The angle of the tibiotarsal joint of the hindleg (*θ*_hTT_) changed similarly to the angle *θ*_hFT_ in both types of jumps (Fig. 4C, D), which accords with previous studies. In case of the attack jump, the angle *θ*_hTT_ changed similarly among insects in the push-off phase, regardless of the initial posture (Fig. 4C). The initial values were in the range of 122 deg to 154 deg. In the preparation phase, the angle *θ*_hTT_ of the estimated curve decreased by about 20 deg from 130 deg. In the push-off phase, the angle *θ*_hTT_ of the estimated curve increased by about 50 deg and reached a plateau. In the case of the escape jump, the angle *θ*_hTT_ was similar to the angle *θ*_hFT_ (Fig. 4D). The initial value of the angle *θ*_hTT_ was in the range of 89 deg to 143 deg. The angle *θ*_hTT_ of the estimated curve decreased slightly (by about 10 deg) in the preparation phase and increased by about 40 deg in the push-off phase.

### Kinematics analysis of forelegs in jumping

Forelegs contributed to supporting the insect’s body and to adjusting its posture during take-off. In locusts, the forelegs control jump trajectory (Santer et al. 2005). To confirm how crickets used their forelegs during attack jumps, we analyzed the kinematics of foreleg joints (Fig. 5). We observed that crickets swung their forelegs backward during attack jumps. To describe this movement, we calculated the angle between the femur and the pronotum in the foreleg (*θ*_fFe_). In the case of the attack jump, the angle *θ*_fFe_ changed in stereotypical patterns, although there were time lags among individuals (Fig. 5A). In the preparation phase, the angle *θ*_fFe_ decreased according to the foreleg’s movement from the front to the side. The initial value was in the range of 61∼165 deg, which was a large variation among individuals. Then, during the push-off phase, the forelegs scraped the ground and moved backward. At that time, the angle *θ*_fFe_ decreased sharply. The estimated curve indicated that the angle *θ*_fFe_ decreased from about 120 deg to 40 deg in 50 ms in jumping (including both preparation and push-off). In the case of the escape jump, the angle *θ*_fFe_ decreased simultaneously with the push-off of the hindleg (Fig. 5B). In the preparation phase, the angle *θ*_fFe_ was initially in the range of 80∼123 deg and decreased slightly. During the push-off phase, the angle *θ*_fFe_ decreased significantly; the estimated curve indicated that the angle *θ*_fFe_ decreased from about 100 deg to 50 deg during the hindleg’s pushing off.

**Fig. 5.**
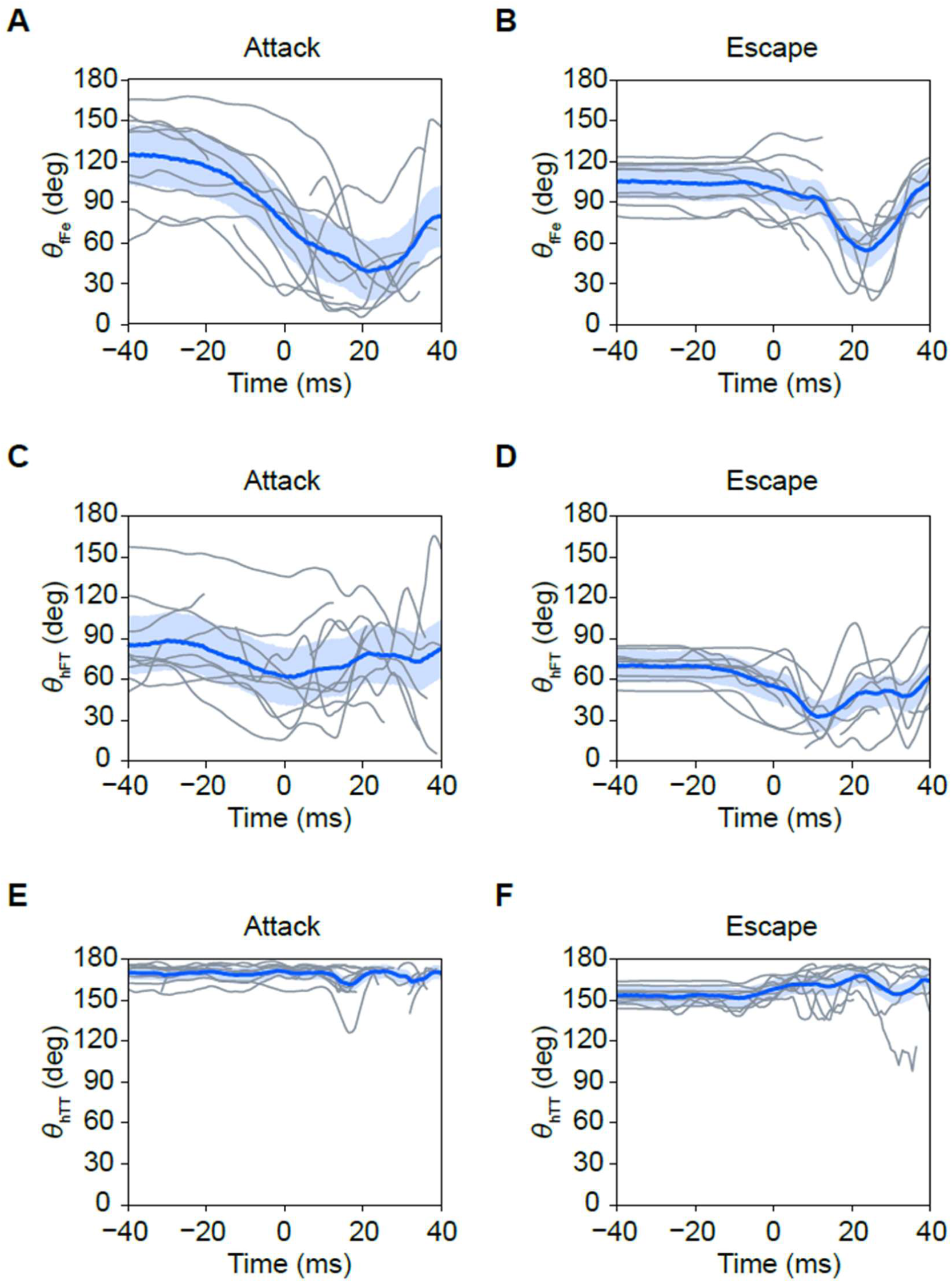
Time validation of the angle of foreleg in attack and escape jumps. (A, B) Time validation of the angle between femur and pronotum in the foreleg (*θ*_fFe_) during attack and escape jumps. (C, D) Time validation of the angle of the femorotibial joint of the foreleg (*θ*_fFT_) in attack and escape jumps. (E, F) Time validation of the angle of the tibiotarsal joint of the foreleg (*θ*_fTT_) in attack and escape jumps. The angles were calculated in each frame using 3D coordinates. All data series were aligned with a time point when crickets start to extend hindlegs as 0 ms. Colored lines indicate the estimated 50th percentile, and the gray area indicates the estimated 25th–75th percentile that was based on the statistical model.

We then analyzed the angle of the femorotibial joint of the foreleg (*θ*_fFT_) because it affected rolling and yawing movements of jumping in locusts (Santer et al., 2005). The angle *θ*_fFT_ in attack jumps varied by individual (Fig. 5C). As an overall trend, the angle *θ*_fFT_ gradually decreased by approximately 45 deg in the preparation phase. Then, it increased sharply by ∼100 deg in the push-off phase. We could not capture the characteristics of the angle *θ*_fFT_ with the statistical model. In the case of the escape jump, the angle *θ*_fFT_ started to decrease at the same time as the hindleg’s preparation (Fig. 5D). Then, it increased at the same time in the push-off phase. The estimated curve indicated that the angle *θ*_fFT_ decreased from 70 deg to 30 deg.

We also analyzed the angle of the tibiotarsal joint of the foreleg (*θ*_fTT_) to examine how firmly crickets stand on the ground with their forelegs. The angle *θ*_fTT_ in attack jumps changed little over time (Fig. 5E). The estimated curve indicated that *θ*_fTT_ in attack jumps ranged between 160 and 170 deg, which was larger than *θ*_fTT_ during escape jumping. This indicated that in the preparation phase, crickets extended their forelegs more than in escape jumping. The angle *θ*_fTT_ in escape jumping increased slightly, from 150 deg to 170 deg, according to the hindleg’s pushing off (Fig. 5F).

### Direction of body axis in jumping

To investigate how the direction of the body axis was determined as a result of the adjustments made by the forelegs, we measured the angle of the longitudinal body axis relative to the ground (*θ*_Axis_) and examined whether it differed between attack and escape jumps (Fig. 6). For simplicity, we used side-view videos made using a single camera to measure the axis angle. In total, 10 jumps were analyzed. In the case of attack jumps, the body angle was 41.0 ± 5.2 deg (mean direction ± angular deviation), while for escape jumps, it was 14.4 ± 9.0 deg. Thus, the body angle during attack jumps was larger than that during escape jumps. To confirm whether the values for these two groups showed the same distribution, we performed the Mardia-Watson-Wheeler test, and the result was *p* = 0.002.

**Fig. 6.**
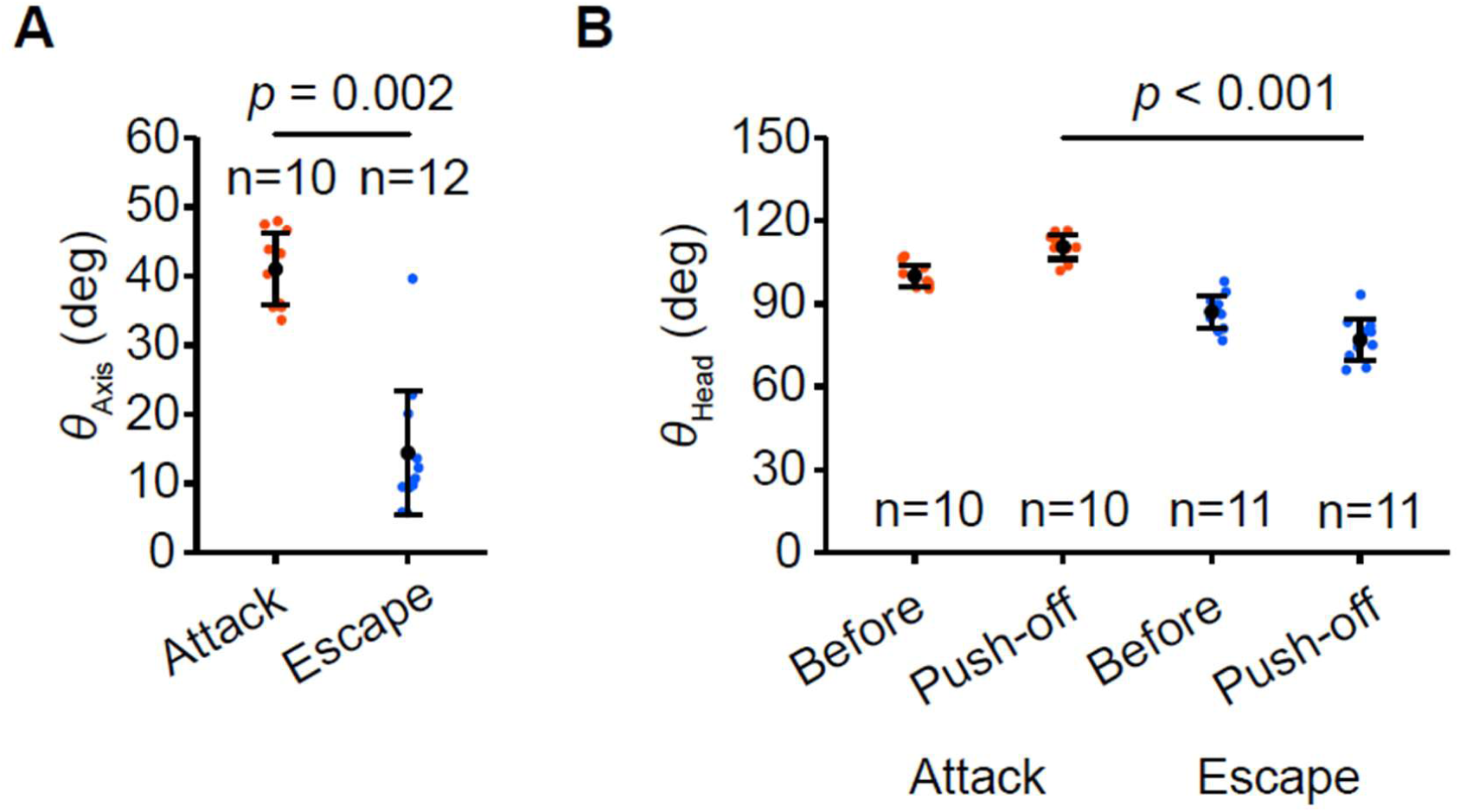
Angle of the axis and the head in attack jumps and escape jumps. (A) Angles of longitudinal body axis relative to the ground. (B) Angles of the front side of the head relative to the longitudinal body axis. Before: before the preparation for the jump. Push-off: when crickets pushed off the ground using their hindlegs. Colored dots represent measured data. Black dots represent the mean angle, and the error bar shows circular standard deviation. The *p-value* was calculated using the Mardia-Watson-Wheeler test.

### Head movement in jumping

Prior to attack jumps, attacking crickets characteristically stuck their mandibles out and hooked them onto the opponent’s mandibles, as mentioned above (Fig. 2). To clarify this motion quantitatively, we measured the angle of the head relative to the body (*θ*_Head_) during the jump and examined whether there was a difference in head angle between attack and escape jumps (Fig. 6B). We measured the head angle both before and during the jump because the crickets moved their head before jumping. In the case of the attack jumps, the head angle was maintained at 100.0 ± 3.9 deg before they jumped (Fig. 6B). During the attack jumps, the head angle was 110.5 ± 4.4 deg. The angle change for each cricket was 10.5 ± 3.4 deg (Fig. 6B). In the case of escape jumping, the head angle was 87.0 ± 5.9 deg before they jumped, and it decreased to 77.1 ± 7.3 deg during the jump (Fig. 6B).

### Overall kinematics during takeoff

To compare the difference in driving force between attack and escape jumps, we calculated the initial velocity and acceleration (Fig. 7). The initial velocity of the attack jumps was 0.50 ± 0.08 m s^−1^ (mean ± SD), which was greater than the initial velocity of the escape jumps, 0.32 ± 0.15 m s^−1^ (*p* = 0.030, Student’s t-test). The acceleration was 25.48 ± 5.23 m s^−2^ for attack jumps and 20.30 ± 10.47 m s^−2^ for escape jumps. There was no significant difference between attack and escape jumps (*p* = 0.302, Student’s t-test).

**Fig. 7.**
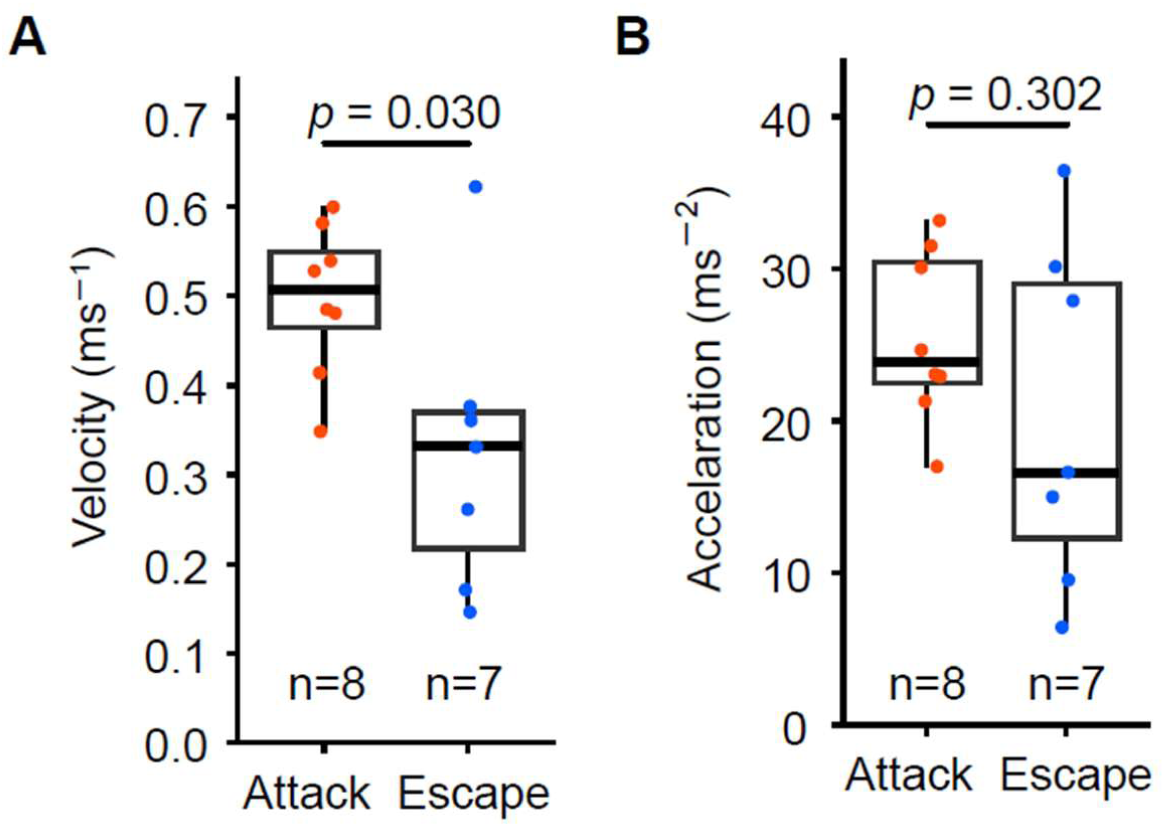
Initial velocity and acceleration of attack and escape jumps. (A) The initial velocity of attack jumps was significantly higher than that of escape jumps (*p* = 0.030, Student’s t-test). (B) There was no significant difference in acceleration between attack jumps and escape jumps (*p* = 0.302, Student’s t-test). Boxplots show the minimum value, median value, maximum value, first quartile, and third quartile. Dots represented individual jumps. The red dots represent attack jumps. The blue dots represent escape jumps. The number of samples is shown at the bottom of the boxplot. The initial velocity and acceleration are calculated from the displacement of the hind legs between the start of the jump and when the hind legs leave the ground.

## Discussion

### Fighting between males

In this study, we show the attack behavior that occurred in crickets jumping to flip over their opponent (Fig. 1, Movie 1). Our observations agree with the motion that previous reports describe as pushing, rushing, or headbutting after antenna fencing (Adamo and Hoy, 1995; Alexander, 1961; Stevenson et al., 2000).

Our results demonstrate that crickets perform this offensive behavior by interlocking their mandibles onto the opponent’s mandibles from below without biting (Fig. 2). In contrast, previous reports described that crickets sometimes bite opponents with their mandibles (Adamo and Hoy, 1995; Judge and Bonanno, 2008; Stevenson et al., 2000). We suggest that crickets primarily employ the attack jump, rather than using biting, to defeat opponents. This type of behavior is more similar to the throwing motion seen in some rhinoceros beetles and stag beetles (Goyens et al., 2015; Hongo, 2003) than to a simple headbutting, as categorized previously (Adamo and Hoy, 1995; Alexander, 1961). Since aggressive behavior is generally associated with territorial disputes, the cricket may apply this motion to try to drive its opponent out of its territory. Additionally, being able to flip an opponent over can demonstrate that the attacker is stronger than their opponent. Even with other insects that do not employ flipping or throwing their opponent, simply pushing their opponent away demonstrates that they are stronger.

### Kinematics of hindlegs in jumping

We gained an understanding of the behavior of crickets utilizing their well-developed hindlegs. The hindlegs are presumed to play a central role in generating enough power to flip an opponent. Therefore, appropriate control of the hindlegs is necessary to accomplish the task. Crickets can perform various movements utilizing their hindlegs even at high speeds. We demonstrated that crickets use their hindlegs differently for attack jumps compared to escape jumps. In attack jumps, contraction of the hindlegs was frequently observed during the preparation phase, suggesting that crickets actively load power prior to take-off. In contrast, such contractions were less commonly observed in escape jumps, likely because rapid reaction to a threat is prioritized over force accumulation. Indeed, crickets have some strategy of escape and use a suitable way depending on the type of stimulus (Sato et al., 2022). In both jumps, crickets started jumping with a similar angle of the hindleg joints. This indicates that the angle is suitable as an initial posture to exert power. Further research on the musculoskeletal system will be required to clarify whether this angle is indeed mechanically optimal.

During the take-off phase of attack jumps, crickets extended their hindlegs more forcefully and sustained the extension briefly after take-off. This suggests that attacking crickets fully extend their hindlegs to maximize jumping performance. In escape jumps, however, crickets immediately transition to running after landing, as continuous locomotion is crucial for successful evasion (Gras and Hörner, 1992; Hiraguchi and Yamaguchi, 2000; Tauber and Camhi, 1995). This usage of the hindlegs enables a smooth transition from jumping to running behavior.

### Kinematics of forelegs

We also gained an understanding of the kinematics of forelegs during the jumps. Forelegs can play a role in adjusting and maintaining the posture for jumping in some insects. Froghopper insects control the azimuth direction in rapid jumping using their foreleg (Santer et al., 2005). The motion of the foreleg in the jumping of crickets has not been described in detail. We showed that the foreleg movement of attacker crickets consists of sequential motions: extend in the preparation phase, stroke the ground, and swing back forward (Fig. 5).

Our data shows that attacker crickets extend forelegs, increasing the angle of the joint between the coxa and the femur (*θ*_fFe_). This could work for controlling the launch orientation of attack jumps. The angle *θ*_fFe_ varied among trials, indicating it could play a role in adjusting the direction of the body axis depending on the posture of the attacked cricket. Indeed, the angle of the body axis in attack jumps was greater than that in escape jumps (Fig. 6A). This is also supported by the posture of crickets when they relax, seen in the initial posture in the escape jump before stimulation, which has less variation (Fig. 6B). On the other hand, a previous study suggested that the cricket *A. domesticus* may control the takeoff angle in voluntary jumps using its hindlegs (Hustert and Baldus, 2010). Although the usage of hindlegs in jumping differs between these species*, G. bimaculatus* shows less abduction by the proximal leg joints of the hindleg than *A. domesticus*. The difference in the musculoskeletal system and its mechanical properties may cause the usage of the appendage.

### Overall kinematics of the attack jump

The kinematic strategy of the attack jump is different from that of the escape jump. To flip the opponent by transmitting the energy efficiently, crickets need to behave in a way different from that of just simple locomotion. In attack jumps, the angle of the body axis relative to the ground was greater than that in escape jumps. This tilt of the body is formed by extension of the forelegs and probably the midlegs, and by putting their abdomen to the ground, as shown before. This posture is considered to detach the attacked crickets from the ground by thrusting them diagonally upwards. The attacking crickets also had another kinematic strategy in the motion of their head: the difference in head motion also plays a role in interlocking attacked crickets with their mandible (Fig. 7). When crickets do not lock anywhere, the point of force application could be misaligned. After adjusting their posture, crickets propulsed their bodies using their hindlegs and thrusted the opponent. In the attack jump, crickets performed at greater speed and similar acceleration relative to that in escape jumps, while pushing the opponent. Overall, it appears that crickets use three strategies to attack opponents: fixing the point of force application, adjusting the direction of jump, and strong propulsion using the hindleg. Further study is needed to determine how effective this strategy is for thrusting the opponent.

## Acknowledgements

A part of this work was supported by Japan Society for the Promotion of Science JSPS Fellowship [Grant-in-Aid for JSPS Fellows, grant number 24KJ1681] to A.M., and KAKENHI [Grant-in-Aid for Scientific Research (A), grant number 22H00216 and 23H00481, and Grant-in-Aid for Challenging Exploratory Research, grant number 22K19795] to H.A.. We thank Dr. Katsushi KAGAYA (Kitami Institute of Technology) for advice on statistics. We also thank Dr. Madoka KITAKAWA (Kobe University) and Dr. Emily M. Standen (University of Ottawa) for their critical reading of our manuscript and their valuable comments. We are grateful to Dr. Elizabeth Nakajima for proofreading the English manuscript.

## Competing interests

The authors declare no competing or financial interests.

## Author contributions

Conceptualization: H.A.; Methodology: A.M, H.A.; Experiment A.M., K.T.; Data analysis: A.M., K.T., H.A.; Resources: H.A.; Writing, review & editing: A.M., K.T., H.A.; Visualization: A.M.; Supervision: H.A.; Project administration: H.A.; Funding acquisition: A.M., H.A.

## Funding

This work was supported by Japan Society for the Promotion of Science [Grant-in-Aid for JSPS Fellows, grant Number 24KJ1681] to A.M., and KAKENHI [Grant-in-Aid for Scientific Research (A), grant number 22H00216 and 23H00481, and Grant-in-Aid for Challenging Exploratory Research, grant number 22K19795].

## Data availability

All relevant data can be found within the article and its supplementary information.

---

**Fig. S1.**
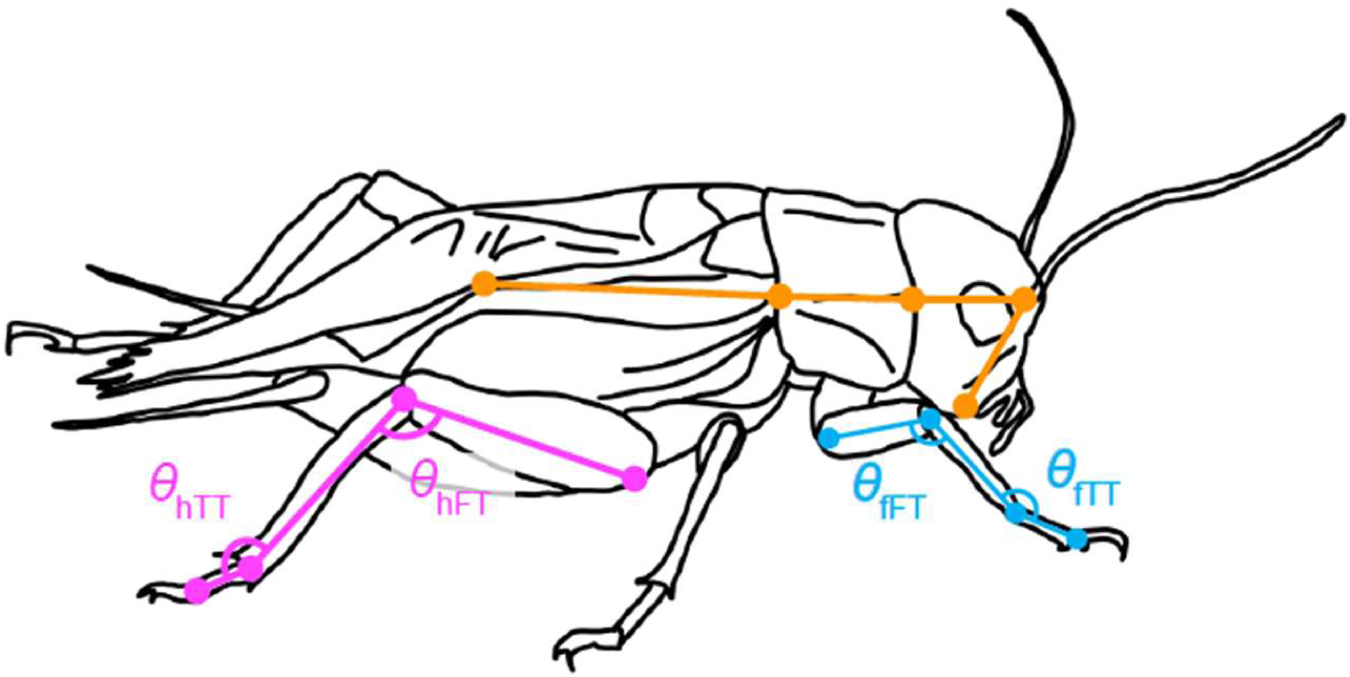
Coarse-grained body of cricket for video tracking Tracking was performed at the points noted in the figure. *θ*_fFT_: Angle of femorotibial joint of foreleg, *θ*_fTT_: Angle of tibiotarsal joint of foreleg. *θ*_hFT_: Angle of femorotibial joint of hindleg, *θ*_hTT_: Angle of tibiotarsal joint of hindleg.

**Suppl. Table 1.**
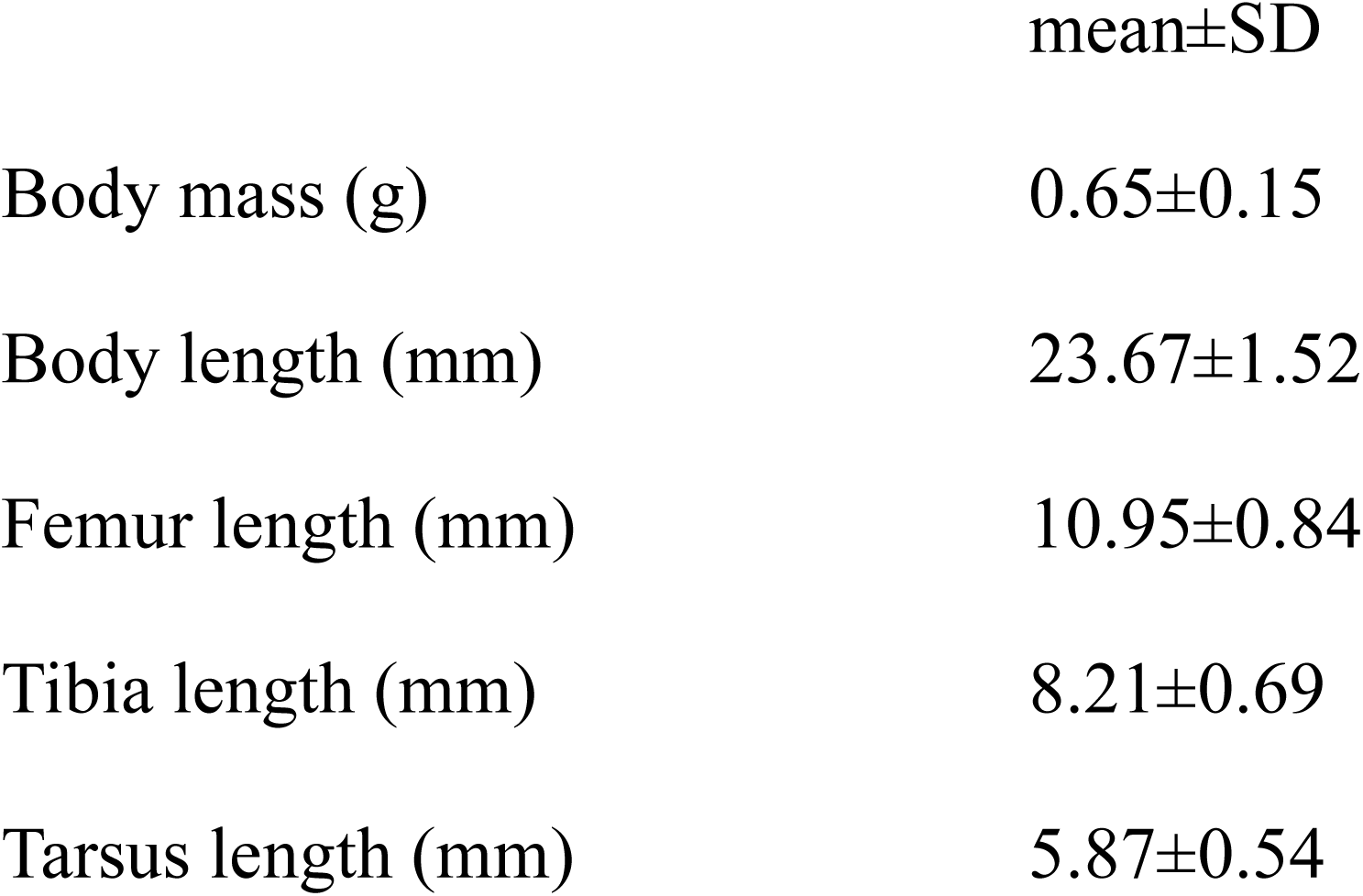
Body size of the crickets (N = 33)

## Notes

### Competing Interest Statement

The authors have declared no competing interest.

