## Supplementary figures and images for "Opponent-flipping tactics in aggressive interactions of the cricket *Gryllus bimaculatus*"

### Supplemental Figure 1

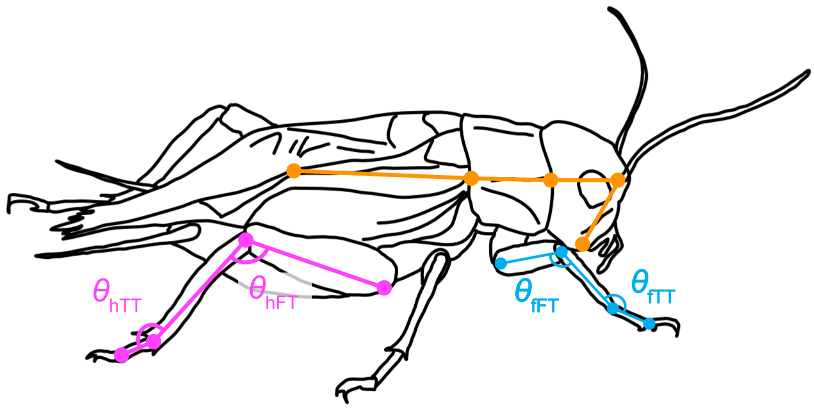


Figure S1
